# Social interactions within the family enhance the capacity for evolutionary change

**DOI:** 10.1101/115014

**Authors:** BJM Jarrett, M Schrader, D Rebar, TM Houslay, RM Kilner

**Affiliations:** Department of Zoology, University of Cambridge, Downing Street, Cambridge, CB2 3EJ; Department of Biology, The University of the South, Sewanee, TN 37383; Centre for Ecology and Conservation, University of Exeter (Penryn Campus), Cornwall, TR10 9FE

## Abstract

Classical models of evolution seldom predict evolution in the wild. One explanation is that the social environment has important, yet overlooked, effects on how traits change in response to natural selection. We tested this idea with selection experiments on burying beetles (*Nicrophorus vespilloides*), sub-social insects that exhibit biparental care. Populations responded to selection for larger adults only when parents cared for their offspring, and responded to selection for smaller adults only when we prevented parents from providing care. Comparative analyses revealed a similar pattern: evolutionary increases in species size within the genus *Nicrophorus* are associated with the obligate provision of care. Synthesising our results with previous studies, we suggest that cooperative social environments enhance the response to selection whereas conflict can prevent further directional selection.

## Main text

Predicting the rate at which populations can evolve and adapt in a rapidly changing world is a major challenge for evolutionary biology^1^. A key problem is to explain how rapidly traits change in response to selection. The breeder’s equation summarizes classical genetic models of evolution by suggesting that the magnitude of evolutionary change in any given trait depends simply on the extent to which that trait contributes to fitness (the strength of selection), and the degree to which it is transmitted to the next generation by genetic variation (the trait’s heritability)^2^. Yet these two parameters are seldom sufficient to predict how evolution will proceed in the wild^3,4^. One suggestion is that this is because the social environment has an additional causal influence on the response to selection^5-9^. An individual’s social environment derives from its interactions with conspecifics. Variation in the social environment can contribute to variation in an individual’s phenotype, much as the abiotic environment does^10,11^. An important difference, though, is that there is genetic variation in the social environment. This means that the social environment can be inherited and can therefore change the response to selection of the traits that it induces^6-9^.

Specifically, mathematical analyses show that when there is a large and positive effect of the social environment on trait expression, it increases a trait’s response to selection and accelerates evolutionary change. But if the effect of the social environment is negative, it prevents any response in the trait to selection and impedes evolutionary change^6-9,12-16^. Previous experiments with domesticated species have supported that latter prediction by showing that competitive interactions can prevent selection for traits of greater economic value to farmers, such as increased body size^13-17^. However, it is unclear whether the social environment can ever causally accelerate trait evolution in animal populations. Nevertheless, theoretical work^6-9^ and correlational analyses of the outcome of natural selection using large pedigreed datasets collected from wild animals, both suggest it is likely^18^.

We tested whether the social environment within the family can promote the evolution of burying beetle size (*Nicrophorus vespilloides*) using experiments on wild-caught individuals. This species exhibits facultative biparental care, which makes it ideal for experimental manipulations of the social environment (e.g. ref. 19). Both parents work together to prepare the carrion nest by removing the fur or feathers from the dead body, rolling the flesh into a ball and burying it underground. Larvae hatch from eggs laid in the soil nearby and crawl to the carcass nest, where they take up residence. There they feed on the flesh themselves, but are also tended by their parents who guard them and transfer resources through regurgitation^20^. However, if parents are removed after nest preparation is complete, but before the larvae hatch, then larvae can complete development without any post-hatching parental care at all^19,21^. After roughly five days, larvae disperse away from the carcass to pupate in the soil.

We focused on the evolution of adult size for three reasons. First, size is strongly associated with fitness in this species^20^. Competition for the carrion breeding resource can be intense, and larger beetles are more likely to win fights for ownership of carcass (e.g. ref. 22). Second, adult size is known from previous work to vary with aspects of the family social environment that larvae experience during development, including social interactions with siblings^23^ and parents^21^. Third, we found that the heritability of adult size is very low. We used techniques from classical quantitative genetics to estimate the heritability of adult size, in environments where parents provided post-hatching care for offspring (hereafter Full Care), and in environments where they provided no post-hatching care, because they were experimentally removed (hereafter No Care). In both environments, the heritability of adult body size did not differ from zero (estimate ± s.e., Full Care: *h^2^* = 0.08 ± 0.12; No Care: *h^2^* = 0.05 ± 0.30, see Supplementary Materials). These estimates are similar to estimates of the heritability of adult size in the congeneric *N.pustulatus*^24^. The breeder’s equation^2^ therefore predicts that body size should exhibit negligible change in response to selection in the short term. This gave us the opportunity to separate the effect of the social environment on the way in which body size responds to selection from effects due to the heritability of body size alone (because the latter should be virtually non-existent).

To test whether the social environment causally influences the response to selection, we carried out an artificial selection experiment on eight laboratory populations (see Methods). Importantly, we varied the social environment among the populations so that we could analyse its causal influence on the response to selection: half the populations experienced Full Care during development (N = 4 populations), the other half had No Care (N = 4 populations). We then exposed half of the populations within each Care environment to selection for increased adult body size (Large), while the remaining populations experienced selection for decreased adult body size (Small, see Methods). Thus we had four types of experimental populations, each replicated twice: Full Care Large, Full Care Small, No Care Large, and No Care Small. We selected on body size for seven generations, generating over 25,000 beetles.

For each experimental treatment, we measured the cumulative selection differential and response to selection, and used these measures to estimate the realised heritability of adult body size (see Methods). This gave us a measure of the extent to which body size could be changed by artificial selection. The breeder’s equation predicts that the realised heritability of body size should not differ among the treatments. However, we found instead that the realised heritability of adult body size varied among the four types of experimental treatments (care × selection × cumulative selection differential: F_3,44_ = 6.87, P < 0.001, Fig. 1). Furthermore, the realised heritability of body size was relatively high, and significantly different from zero, for the Full Care Large treatment (0.090 ± 0.021), where mean body size increased across the generations, and for the No Care Small treatment (0.105 ± 0.033), where mean body size correspondingly decreased. For these two treatments we therefore conclude that the social environment during development enhanced the capacity for evolutionary change in adult body size, and to a similar degree whether selection was for increased or decreased body size.

**Figure 1.**
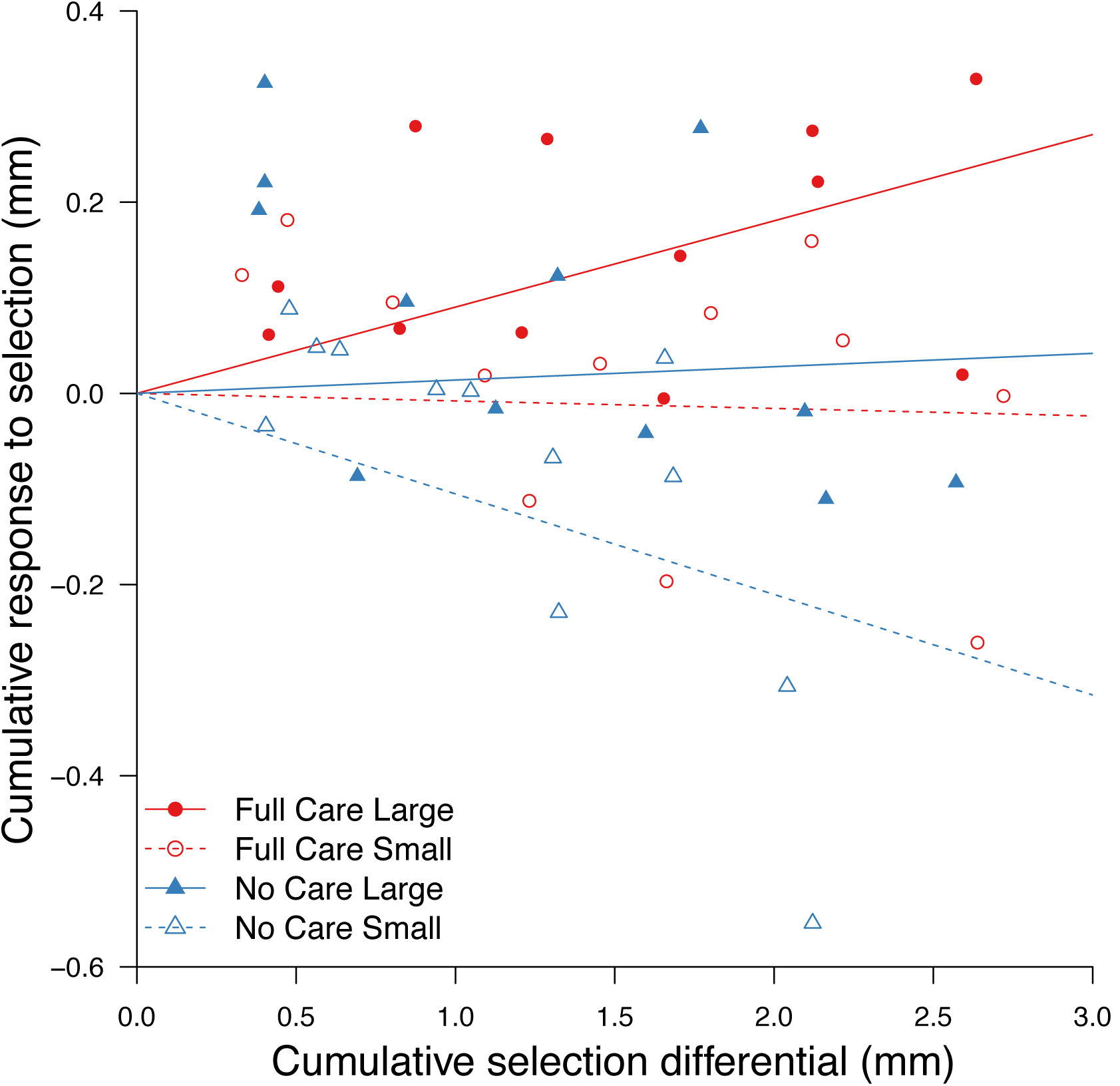
The realised heritability of body size, as a function of the different selection regimes and social environments. The realised heritability is given by the regression slopes, forced through the intercept. For each treatment the gradient of these regression lines ± S.E are: Full Care Large, 0.090 ± 0.021; Full Care Small, –0.008 ± 0.023; No Care Large, 0.014 ± 0.033; No Care Small, –0.106 ± 0.033. The cumulative selection differential is the difference between the population mean and the mean of the retained subset of the population. This is summed across the seven generations. The cumulative response to selection is the difference between the mean of the population and the mean of the population in the subsequent generation, and is also summed. The two replicates for each treatment were pooled for the regression, as they did not differ (see Supplementary Materials).

By contrast, in the Full Care Small and the No Care Large treatments, the realised heritability of adult body size was not significantly different from zero (Full Care Small: −0.008 ± 0.023; No Care Large: 0.014 ± 0.033). Mean adult body size did not change over the course of the selection experiment for individuals from either of these treatments (Fig. 1).

The next step was to determine how the two contrasting social environments in our selection experiment could influence evolutionary change in adult size. Previous work has shown that the mass a larva attains by the time it disperses away from the carcass strongly influences the size of the adult that then emerges^25^. Furthermore, larval mass at dispersal depends on the number of larvae competing during development for the finite resources on a carcass^23^. Building on these results, we identified three social factors that influence larval mass at dispersal. The first is clutch size, because it influences the number of larvae competing for carrion. However, it is not the sole determinant of brood size on a carcass. Larger females lay a larger clutch^26^ but have fewer surviving larvae that disperse from the carcass (see Methods, Supplementary Fig. 1), presumably due to a greater incidence of filial cannibalism^27^. Brood size at dispersal is therefore different from clutch size, and is the second factor influencing larval mass at dispersal. The third factor is the presence or absence of parents after hatching. This factor is important because it influences the relationship between brood size and larval size at dispersal, especially for broods of 10 or fewer larvae. When parents are present, and there are only a few larvae on the carcass, each consumes more carrion and is larger at dispersal^23^. However, when parents are absent, each larva typically attains only a low mass by the time it disperses to pupate, because larvae seemingly help each other to colonize and consume the carcass^23^. Thus larvae in small broods cannot attain a large mass at dispersal when parents are absent, but they can when parents are present.

We suggest that selection on these three elements of the social environment combined to cause correlated change in body size in the Full Care Large lines and the No Care Small lines (see Supplementary Materials). In the Full Care Large treatment (Fig. 2a), we selected for larger adults. They produced larger clutches (Supplementary Fig. 2), but produced fewer (Supplementary Fig. 3) and therefore larger dispersing larvae (presumably due to greater levels of filial cannibalism). They matured into larger adults themselves. Likewise, in the No Care Small treatment (Fig. 2b) we selected for smaller adults and they laid a smaller clutch (Supplementary Fig. 2). Since these broods developed without parents, the resulting smaller broods yielded smaller larvae (Supplementary Fig. 3), which matured into smaller adults. In each treatment, we effectively selected a social environment on the carcass that induced the production of more individuals with either a larger (Full Care Large) or smaller (No Care Small) body size. Furthermore, these selected individuals then produced a similar social environment for their offspring. This explains why these lines responded to selection on body size, despite the very low heritability of body size.

**Figure 2.**
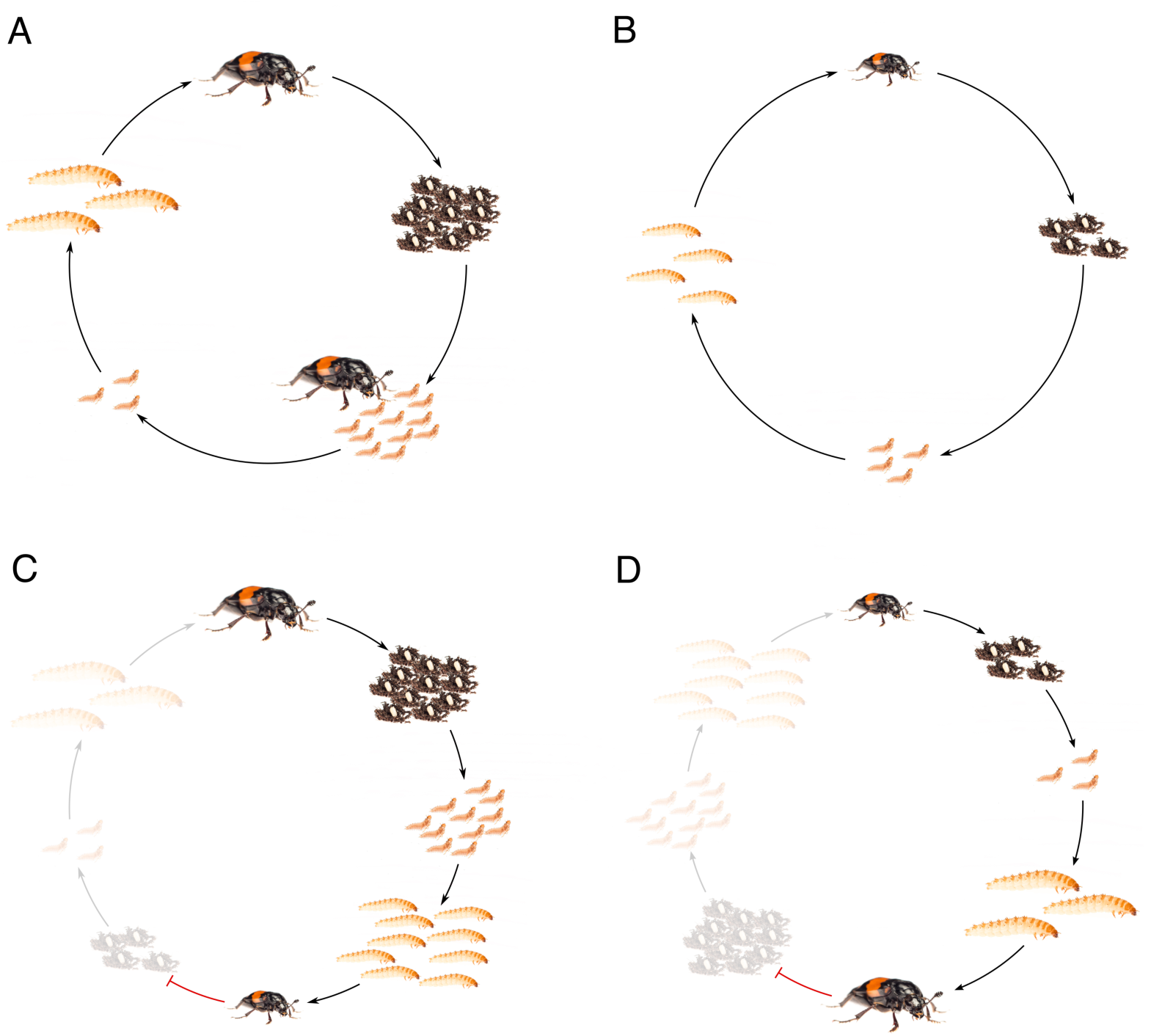
The effect of the social environment on the response to selection, in each of the experimental treatments. (A) and (B) show how the social environment enhances the capacity for evolutionary change; (C) and (D) show how the social environment could prevent evolutionary change. (A) Full Care Large: large beetles lay many eggs, but are more likely to cannibalize larvae and so have relatively small broods that yield large larvae, which mature in large adults. (B) No Care Small: small beetles lay fewer eggs, which yield a small brood of small larvae that mature into small adults. (C) No Care Large: large beetles lay many eggs, which yield a larger brood of small larvae that mature into small adults and are selected out of the experimental population; and (D) Full Care Small: small beetles lay fewer eggs which yield a small brood of large larvae that mature into small adults and are selected out of the experimental population.

We observed very little change in body size in the other experimental populations (No Care Large, Full Care Small). This was predicted by the classical estimates of heritability, but it might also be attributed to effects of the social environment, which could have cancelled out the effects of selection at each generation (see ref. 28). For example, in the No Care Large treatment (Fig. 2c), selecting for larger adults yielded smaller individuals in the next generation. The larger adults laid a larger clutch (Supplementary Fig. 2), but with no parents present after hatching to cannibalize offspring, these larger clutches yielded relatively large broods (Supplementary Fig. 3) of smaller larvae, which matured into smaller adults. Similarly, in the Full Care Small treatment (Fig. 2d) selection for smaller adults yielded larger adults in the following generation. The smaller adults laid a smaller clutch (Supplementary Fig. 2), which in turn yielded a smaller brood (Supplementary Fig. 3) of relatively large larvae that matured into large adults.

We explicitly tested the conclusions set out in Fig. 2, by comparing the slope of the regression between dam size and progeny size (see Supplementary Materials). Fig. 2a and 2b predict that in the Full Care Large and No Care Small treatment, this correlation should be positive, whereas Fig. 2c and Fig. 2d predict it should be negative in the No Care Large and Full Care Small treatment. We found that the slopes of these correlations differed significantly among treatments (care × selection × dam pronotum: χ^2^_1_ = 4.13, P = 0.042). The slopes were positive in the Full Care Large (0.134 ± 0.090) and No Care Small treatments (0.094 ± 0.079). However, although they were negative in the Full Care Small treatment (−0.059 ± 0.064), as we predicted, they were positive in the No Care Large treatment (0.117 ± 0.098), which we did not predict.

Our experiments thus find no clear evidence to support the suggestion that the social environment within the family alone prevented evolutionary change in the Full Care Small and No Care Large treatments. They do, however, show that social interactions within the family enhanced the response to selection in the Full Care Large and No Care Small treatment. More specifically, our experiments indicate that parental care is essential to promote a rapid evolutionary increase in body size in *N. vespilloides.*

We tested the merits of this conclusion in a final comparative analysis across the *Nicrophorus* genus, to link our experimental results back to the natural world (see Methods). Different species of burying beetle are remarkably alike in their ecology and appearance^29^. They differ principally in their relative size and in the extent to which parental care is essential for larval growth and survival^30^. Observations of natural burying beetle populations show that adult size is correlated with variation in the size of carrion used by different species for reproduction^20^. Variation in adult body size is correlated with the partitioning of the carrion niche by sympatric species, and enables larger species to favor larger carrion and smaller species to breed on smaller carcasses^20^. We mapped the changes in adult body size across the *Nicrophorus* genus by measuring museum specimens of 49 of the 68 extant species^29^ and placing them on a recent molecular phylogeny of the genus (Fig. 3)^30^. We found that there is considerable variation in body size across the phylogeny, with multiple shifts to both larger and smaller species relative to the ancestral phenotype (Fig. 3). Consistent with our experimental results, we also found that the evolution of very large burying beetles is associated with obligate provision of parental care (PGLS: estimate = 1.57 ± 0.66, t_12_ = 2.40, P = 0.035).

**Figure 3.**
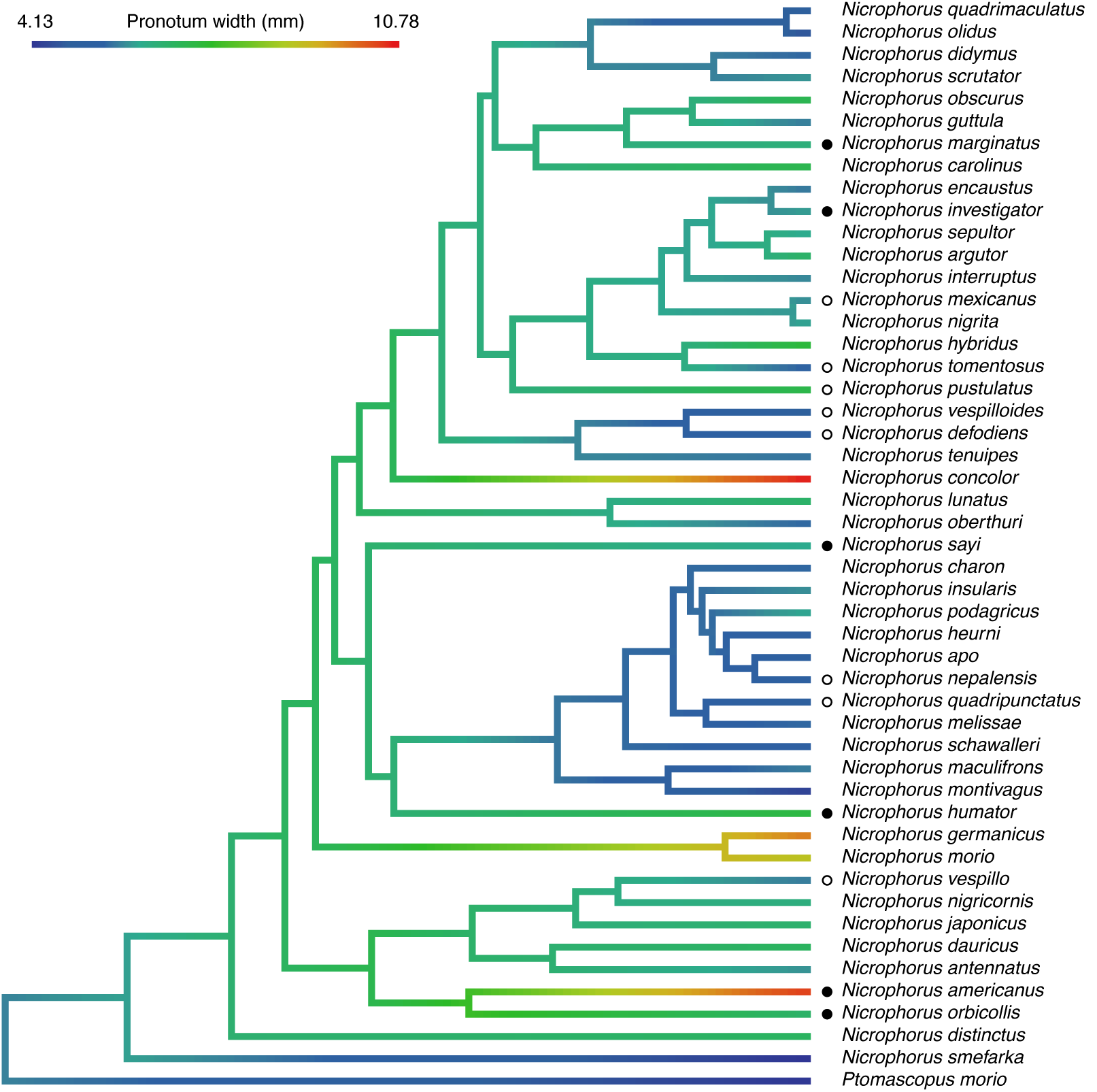
Adult pronotum width of burying beetle species mapped on an existing molecular phylogeny^31^. Black circles indicate species with obligate post-hatching parental care; open circles indicate facultative post-hatching parental care. Species with missing data for parental care have no symbols. Body size data can be found in Supplementary Table 2. Information regarding parental care can be found in Supplementary Table 3.

We conclude that the way in which the social environment influences a trait’s response to selection depends on whether it is associated with social interactions that are cooperative or promote conflict (see ref. 10 for formal definitions of these terms). Previous studies have shown that selection for increased size or productivity also selects for increased aggression. Increased aggression reduces fitness so much that any effects of selection on size cannot be transmitted to the next generation and this prevents evolutionary change^13,17^. This suggests that traits associated with social environments that induce conflict have limited capacity for further directional evolutionary change. Previous work has also demonstrated that, under these conditions, the only way in which increased productivity or size can be artificially selected is by imposing multilevel, group or kin selection^12,13^. That is, a response to selection can be restored only when an explicitly cooperative social environment is artificially created at the same time^32^. Our experiment provides more direct evidence that cooperative interactions enhance the response to selection, and can do so even when selection acts on individuals. In the Full Care Large treatment, selection for increased body size was possible because parents helped small broods of larvae to attain a large size at dispersal. Likewise, in the No Care Small treatment (Fig. 2) selection for decreased body size was possible because cooperative interactions among larvae influence body size^23^: in this case, selection for smaller individuals decreased brood size and the fewer remaining larvae were increasingly unable to help each other grow large. In short, cooperative interactions reinforced selection by magnifying changes in body size across generations, so enhancing the capacity for evolutionary change. Our general conclusion is that the response to selection is likely to be reduced when trait expression is associated with conflict, but enhanced for traits whose expression is associated with more cooperative social environments. Proper characterization of the social environment in which traits are expressed is therefore important not only for understanding a trait’s current adaptive value^10^ but also for predicting its future capacity to evolve and adapt.

## Acknowledgments

This project was funded by a European Research Council grant (310785_Baldwinian_Beetles), and a Royal Society Wolfson Research Merit Award, both to RMK. We are very grateful to S.-J. Sun and D. Howard for providing unpublished information about other burying beetle species, and to C. Creighton for discussion. We thank S. Castañón for help with Matlab; M. Barclay and R. Booth from the Natural History Museum, London for their help with the beetle collections; and K. MacLeod and P. Lawrence for commenting on earlier drafts. A. Backhouse, S. Aspinall and C. Swannack maintained the beetles while A. Attisano, E. Briolat, A. Duarte and O. de Gasperin helped in the laboratory.

## Materials and Methods

The burying beetle genus *Nicrophorus* is distributed primarily throughout the temperate regions of the Northern Hemisphere^29^. So far as is known, the natural history and reproductive biology of all *Nicrophorus* species are broadly similar^20,29,31^ and centre on the use of small carrion as a breeding resource^20^. Although the two other extant genera in the Nicrophorinae also use carrion for reproduction, they lack the elaborate parental care exhibited by *Nicrophorus* species and the associated social interactions that it generates^31,33^. These genera are also less speciose than *Nicrophorus:* there are 68 known species in *Nicrophorus,* one in *Eonecrophorus* and three in *Ptomascopus^29^.* This suggests there is a correlation between the social environment during development and the capacity for diversification in each of these lineages.

## Estimating the heritability of body size in *N. vespilloides*

### *Cultivating* N. vespilloides *in the lab*

All the individuals used in this experiment belonged to a captive colony (kept at a constant temperature: 21°C, with a 16h:8h light:dark cycle) established at the University of Cambridge in 2013 from wild-caught adults collected under licence from local field sites at Byron’s Pool and Wicken Fen in Cambridgeshire, U. K. Adults were housed individually in plastic boxes (12 × 8 × 2cm) filled with moist soil (Miracle Grow) and fed twice a week with ~0.3g of minced beef. For breeding, pairs of unrelated individuals were placed into larger plastic boxes (17 × 12 × 6cm) half-filled with moist soil, provided with a 8–13g freshly thawed mouse carcass and kept in the dark to simulate natural underground conditions. The larvae disperse from the carcass to pupate roughly eight days after pairing. Dispersing larvae were transferred into population boxes (10 × 10 × 2cm), each subdivided into equal cells of 2 × 2 × 2cm and filled with soil. Once pupation was complete (approximately 3 weeks after dispersal), each sexually immature adult was moved to its own individual, uniquely labeled box. Sexual maturity is reached approximately two weeks after eclosion, and beetles were paired for reproduction at this time. No siblings or cousins were paired for breeding.

### Methods

We performed a full-sib/half-sib quantitative genetics experiment to estimate the heritability of body size in *N. vespilloides.* We used two populations of beetles for this experiment, both maintained under the same conditions as stock populations (Full Care) for 11 generations without any selection for body size. Four females were mated to a single male and then each female was given a recently defrosted mouse (10-12g) to breed upon. Once the carcass had been prepared and all eggs laid, approximately 53h after providing the mouse^34^, the female and carcass were removed. The female was placed in a new breeding box and provided with a fully prepared carcass from a donor female. At that time we also prepared an equal number of breeding boxes with just a donor-prepared carcass and no female. The breeding box where the female laid her eggs was checked three times a day for larval hatching. Once larvae started hatching, the larvae were transferred to either the carcass with their mother (Full Care) or to the other carcass without an adult (No Care). Larvae were added until a maximum of 12 larvae were present on each carcass, resulting in mean (± s.e.) brood sizes of 7.85 ± 0.25 in the Full Care, and 8.21 ± 0.24 in the No care environments.

We checked breeding boxes three times daily, and determined that the larvae were ready to disperse when two or more larvae were seen crawling away from the remains of the carcass^24^. At this point the contents of the breeding box were removed and the larvae were counted and weighed individually. The larvae were then placed into individual cells within an eclosion box in the order in which they were weighed so we could relate larval mass to adult size. After eclosion, we anaesthetized the adults with CO_2_. Once anaesthetized, each individual was placed flat under a Canon DSLR camera and photographed. The camera was attached to a stand to ensue consistency in the images obtained and connected to a computer for automatic image labeling. All photographs contained a scale against which the pronotum width of each individual was measured using a custom MatLab script. No statistical methods were used to predetermine sample size.

We analyzed data for each care regime separately, using the package ASreml-R 3.0^35^ in R version 3.3.0^36^. Models included a fixed effect of the number of larvae surviving per brood (mean-centered), a random effect of brood ID to estimate variance due to permanent environmental (including maternal) effects, and a random effect of the pedigree term to estimate the additive genetic variance. (We were unable to partition variance due to maternal effects from that of the permanent environment because no females had multiple broods within a single environment). We then tested the significance of the additive genetic variance in adult size by comparing models with and without the pedigree term using a likelihood ratio test. We estimated χ^2^_nDF_ as twice the difference in model log likelihoods; given that we were testing the effect of a single variance component (nDF = 1), we assumed that the test statistic was asymptotically distributed as an equal mix of χ^2^_0_ and χ^2^_1_ (ref. 37). The heritability of adult size was calculated as V_A_ / V_P_ where V_P_ is the sum of the variance components (additive genetic, permanent environment, and residual) from the model, having conditioned on the fixed effects. We used Wald *F*-tests to estimate the significance of fixed effects.

### Results

The experiment yielded 186 maternal full-sib families and 56 paternal half-sib families in the Full Care environment, and 84 maternal full-sib families and 22 paternal half-sib families in the No Care environment. Mean (± s.e.) brood size in the Full Care was 7.69 ± 0.24 and 5.31 ± 0.30 in the No Care.

We found no evidence for significant additive genetic variance in adult size in either the Full Care (V_A_ = 0.013 ± 0.021, χ^2^_0,1_ = 0.46, P = 0.25) or No Care (V_A_ = 0.008 ± 0.045, χ^2^_0,1_ = 0.03, P = 0.43, Supplementary Table 1) environments. The heritability estimates of adult size were correspondingly close to zero, with large standard errors (h^2^_Full_ = 0.08 ± 0.12; h^2^_No_ = 0.05 ± 0.30). Permanent environment effects (ie effects of the Care treatment and brood size) explained a significant amount of the total phenotypic variation in adult size (conditional on fixed effects) in both Full Care (V_PE_ = 0.046 ± 0.012, χ^2^_01_ = 16.22, P < 0.001; proportion of total phenotypic variance conditional on fixed effects = 0.263 ± 0.065) and No Care (V_PE_ = 0.054 ± 0.025, χ^2^_0,1_ = 6.05, P = 0.007; proportion = 0.361 ± 0.157) environments. For completeness, we ran the same models without any fixed effects (see ref. 38), but this had no meaningful effect on our results.

## Selection experiment

One way to analyse the effect of the social environment on the response to selection is to use cross-fostering to partition out sources of variance in body size to direct, sib-social, or maternal effects^39-41^ and thereby deduce the underlying genetic architecture. However, the downside of this approach is that it requires detailed knowledge of precisely how the social environment influences trait expression: if one key element is overlooked then the analyses are too incomplete to be able to predict the response to selection with any accuracy. For this reason, we chose instead to use an artificial selection experiment. We manipulated the social environment, imposed selection and measured the response. In this way we could confidently attribute any change in the response to selection to our manipulations of the social environment, without making any *a priori* assumptions about which particular aspects of the social environment were important in influencing trait expression.

All the individuals used in the selection experiment belonged to a captive colony established at Cambridge University in 2013 from wild caught adults collected under licence from local field sites at Byron’s Pool and Wicken Fen in Cambridgeshire, U.K. Full details of the protocols used are given in (ref. 19).

### Methods

From the genetically diverse founding population, we started eight populations consisting of four treatments with two replicates per treatment, randomly allocating individuals to treatments. We had two treatments, Provision of Care and Selection for Size, resulting in a 2 × 2 factorial experiment. Provision of Care was manipulated by either leaving or removing both parents 53 hours after pairing, after carcass preparation and egg laying were complete^34^, resulting in a Full Care treatment, and a No Care treatment, respectively. We then imposed two selection regimes on the Full Care and No Care populations: Large and Small. We selected the largest third of the population with the Large regime, and the smallest third of the population under the Small regime. Selection was imposed at the population level and not at the family level. Once the population had been selected, individuals were paired haphazardly, although we ensured cousins and siblings did not breed. All beetles were maintained under the conditions described above. Each population was maintained with at least 25 families per generation, by breeding 40 pairs of beetles for the Full Care populations and 60 pairs for the No Care populations. When it became impossible to sustain populations of this size, the experiment ceased. (We bred extra pairs in the No Care population to ensure there were enough successful families: failure rates are high when initially removing parental care).

At eclosion members of the same sex from each family were temporarily housed in a box together and anaesthetised with CO_2_. Once anaesthetized, each individual was photographed and the body size measured in the same method as described above. Each individual was given a unique ID that we used to identify individuals that were retained to breed in the next generation.

To estimate the potential for evolutionary change in body size in each population, we calculated the realised heritability of body size, as the slope of the regression of the cumulative response to selection against the cumulative strength of selection^42^. Post-hoc pairwise comparisons were adjusted for multiple testing^43^. No statistical methods were used to predetermine sample size.

### Results

The realised heritability did not differ significantly between replicate populations for each treatment (F_40_ = 2.08, P = 0.10). Replicates were therefore pooled for all subsequent analyses. After running the global model, we used pairwise comparisons to compare measures of realised heritability across the different treatments. The Full Care Large and Full Care Small treatments significantly differed from one another in realised heritability (F_22_ = 9.90, P_adj_ = 0.015), as did the Full Care Large and No Care Small (F_22_ = 26.44, P_adj_ = 0.006). There was marginal support for a difference in realised heritability between Full Care Large and No Care Large (F_22_ = 3.95, P_adj_ = 0.072). Realised heritability in the No Care Small treatment differed significantly from that in the Full Care Small (F_22_ = 5.92, P_adj_ = 0.03) and the No Care Large populations (F_22_ = 6.36, P_adj_ = 0.03). The Full Care Small and No Care Large did not differ from one another in their realised heritability (F_22_ = 0.30, P_adj_ = 0.59). Realised heritability estimates for each population are in Supplementary Table 2.

## The effects of the social environment on adult size

The social environment that larvae experience during development influences the size the larvae attain by the time they disperse from the carcass and this, in turn, is strongly correlated with adult size^25^. Three factors contribute to this social environment (see main text): clutch size, brood size at dispersal and the presence (or absence) of parents during larval development^23^. To understand how these different elements of the social environment might have caused the outcome of the selection experiment, we began by investigating how clutch size and brood size are related to adult size.

### a) Relationship between female size and clutch size, or brood size at dispersal

To assess the effect of female size on clutch size we analysed data from^26^ where we manipulated female size experimentally and destructively counted the total clutch size for a breeding attempt after 53 hours when egg laying has ceased^34^. Brood size data were taken from a stock population maintained in the laboratory under the same conditions as the Full Care populations, and assayed when the selected populations were in generation five. Brood size was measured at the point of larval dispersal away from the carcass. Both clutch size and brood size were analysed with a Poisson distribution and a log link function with female size and carcass mass fitted as covariates.

We found that clutch size increased with female size even when accounting for carcass mass (t = 3.63, P = 0.001), whereas brood size at dispersal decreased with female size (t = –2.06, P = 0.04, Supplementary Fig. 1).

The next step was to relate these effects of the social environment to the results of our selection experiment. If the outcome of the selection experiment is attributable to different elements of the social environment, then we predict we should see divergence in clutch size, and brood size at dispersal among the different experimental treatments.

### b) Measurement of clutch size in the experimentally selected populations

Based on the results in Supplementary Fig. 1, we predict that clutch size should be greater in populations where adults are selected to be larger (i.e. Full Care Large and No Care Large) than in populations where adults are selected to be smaller (i.e. Full Care Small and No Care Small). To test this prediction, we estimated clutch size in all eight populations at generation five by counting the number of eggs visible on the bottom of the breeding box. We know from previous work that this measure is strongly correlated with total clutch size^26^. We analysed estimated clutch size using a generalised linear model with a Poisson error structure, and log link function. We included carcass size as a covariate.

As predicted, we found that clutch size in generation five of the selection experiment was greater in the Large selected lines than in the Small selected lines (z = –7.53, P < 0.001), independent of the parental care treatment (z = 1.32, P = 0.19, Supplementary Fig. 2). There was no interaction between selection regime and parental care on clutch size (z = –0.38, P = 0.70).

### c) Measurement of brood size in the experimentally selected populations

We predicted that brood size at larval dispersal should also differ among the experimental populations. Specifically, based on the results in Supplementary Fig. 1, we predicted that members of the Full Care Large populations should have a smaller brood size than members of the Full Care Small populations. In addition, since there is no possibility of filial cannibalism in the No Care populations, we predicted that in these populations brood size should vary in the same way as clutch size, and therefore should be greater in the No Care Large populations than in the No Care Small populations. We measured brood size at larval dispersal in Generation 7 of the selection experiment and pooled both replicates. We analysed estimated brood size using a generalised linear model with a Poisson error structure, and log link function, and tested our prediction by searching for a significant interaction between parental care (Full Care, No Care) and selection regime (Large, Small) on brood size at dispersal. We included carcass size as a covariate.

As predicted, we found a significant interaction between parental care and selection regime on brood size at larval dispersal in generation seven (z = –4.89, P < 0.001). Full Care Large populations had fewer offspring at dispersal than the Full Care Small populations, whereas No Care Large populations had more offspring at dispersal than No Care Small populations (Supplementary Fig. 3).

### d) Testing predictions from Figure 3

From Fig. 3, we predicted that the slopes of offspring size regressed against dam size would differ among the experimental treatments. Specifically, we predicted that the slope would be positive for the Full Care Large and No Care Small lines, because these were the lines in which we observed phenotypic change. And we predicted that the slope would be negative in the No Care Large and Full Care Small lines. We took all the data from all the lines and combined both replicates per treatment for the seven generations of the experiment.

We used R^36^ and the package lme4^44^ to a run a linear mixed model, where we ran a model coding the three-way interaction of Care treatment (Full Care or No Care), selection regime (Large or Small) and dam pronotum width. Also included in the model was carcass size and generation. Dam ID was fit as a random term. Significance was determined by removing the three-way interaction from the model and comparing the output with the full model. The slopes for each experimental treatment were obtained in the same way, but with the appropriate subset of the data for each experimental treatment.

## Phylogenetic analysis of body size

We collected data on *Nicrophorus* body size using the beetle collections at the Natural History Museum in London. We took standardized photographs of representatives from all the *Nicrophorus* species included in a recently published molecular phylogeny^31^, with a constant distance between subject and camera, and including a scale-bar in each picture. There was no sexual size dimorphism in our dataset (t =−1.453, P = 0.15). Therefore body size data from both sexes were pooled for each species. We used the standard practice of quantifying body size by measuring pronotum width, and used a MatLab script to calibrate photographic measurements of pronotum width with the scale bar in each image, using the same method for both experiments detailed above. The full datasets can be found in Supplementary Table 3. Post-hatching parental care was classified as ‘facultative’ or ‘obligate’ using data from the published literature and from personal communication with other burying beetle researchers (N = 14 species, Supplementary Table 4). ‘Obligate’ parental care was defined as the failure of larvae to survive to the third instar when parents were removed.

We used a phylogenetic generalised least squares regression (PGLS) to analyse the relationship between body size and parental care using R version 3.3.0^36^ with packages ape^45^, picante^46^ and caper^47^. Care was coded with a dummy variable that was treated as a factor in (1 = obligate post-hatching parental care, 0 = facultative posthatching parental care). Species without a parental care classification were coded NA.

We removed data obtained through personal communication systematically and repeated the analysis to check whether these data affected our conclusions. They did not. We removed *N. americanus* (est = 0.88 ± 0.35, t_11_ = 2.54, P = 0.028), *N. marginatus* (est = 1.72 ± 0.72, t_11_ = 2.40, P = 0.035), and *N. nepalensis* (est = 1.52 ± 0.71, t_11_ = 2.13, P = 0.056) from our analysis separately, and without all three species (est = 0.85 ± 0.42, t_9_ = 2.05, P = 0.07). The results without *N. nepalensis,* and without all three species, were still marginally significant. More importantly, a large effect size in the same direction was retained: that is, larger species have obligate care (see Main Text).

## Supplementary Figures and Figure Captions

**Supplementary Fig 1.**
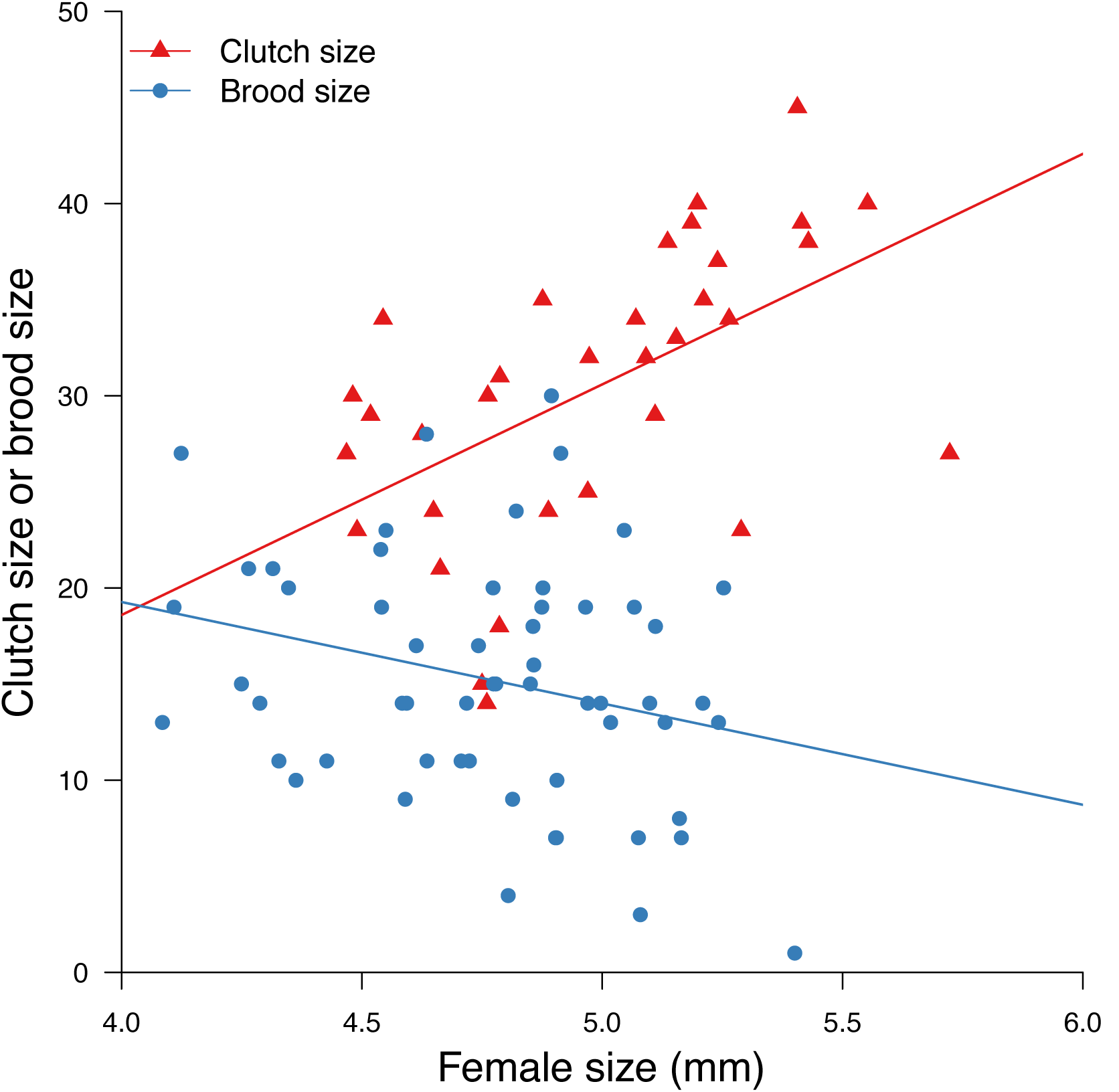
The relationship between female size and a) clutch size (in red triangles); and b) brood size (in blue circles). Clutch size (red line, N = 33) increases with female size. Data were taken from^26^. Brood size (blue line, N = 55), decreases with female size. Female size refers to pronotum width. Each datapoint corresponds to a different female.

**Supplementray Fig 2.**
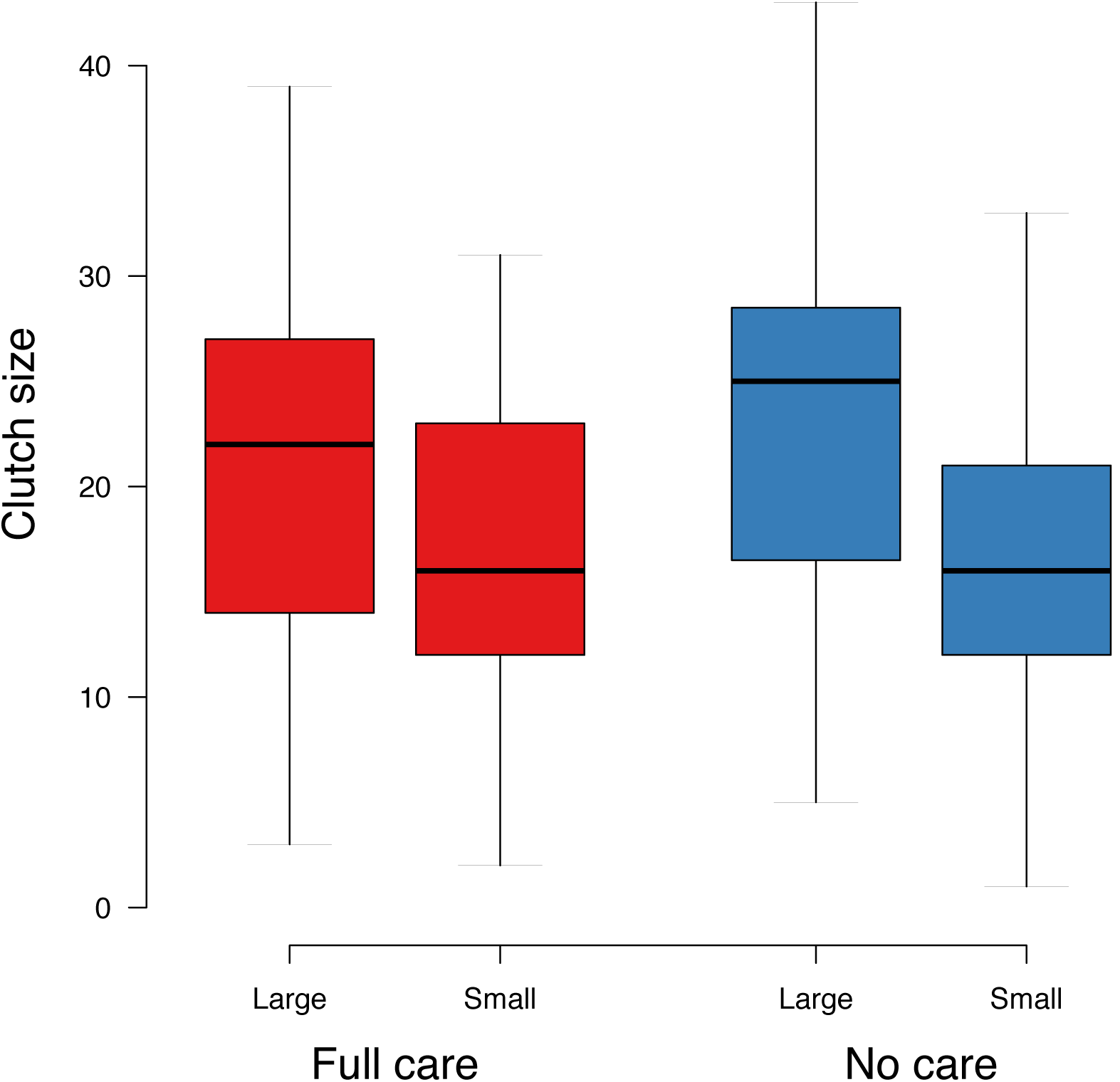
Clutch size at generation five in the four different experimental treatments in the selection experiment: Full Care Large (N=38) and Full Care Small (N=39) (in red); No Care Large (N=51) and No Care Small (N=44) (in blue). Both replicates per treatment are combined. Box plots show median and interquartile ranges.

**Supplementray Fig 3.**
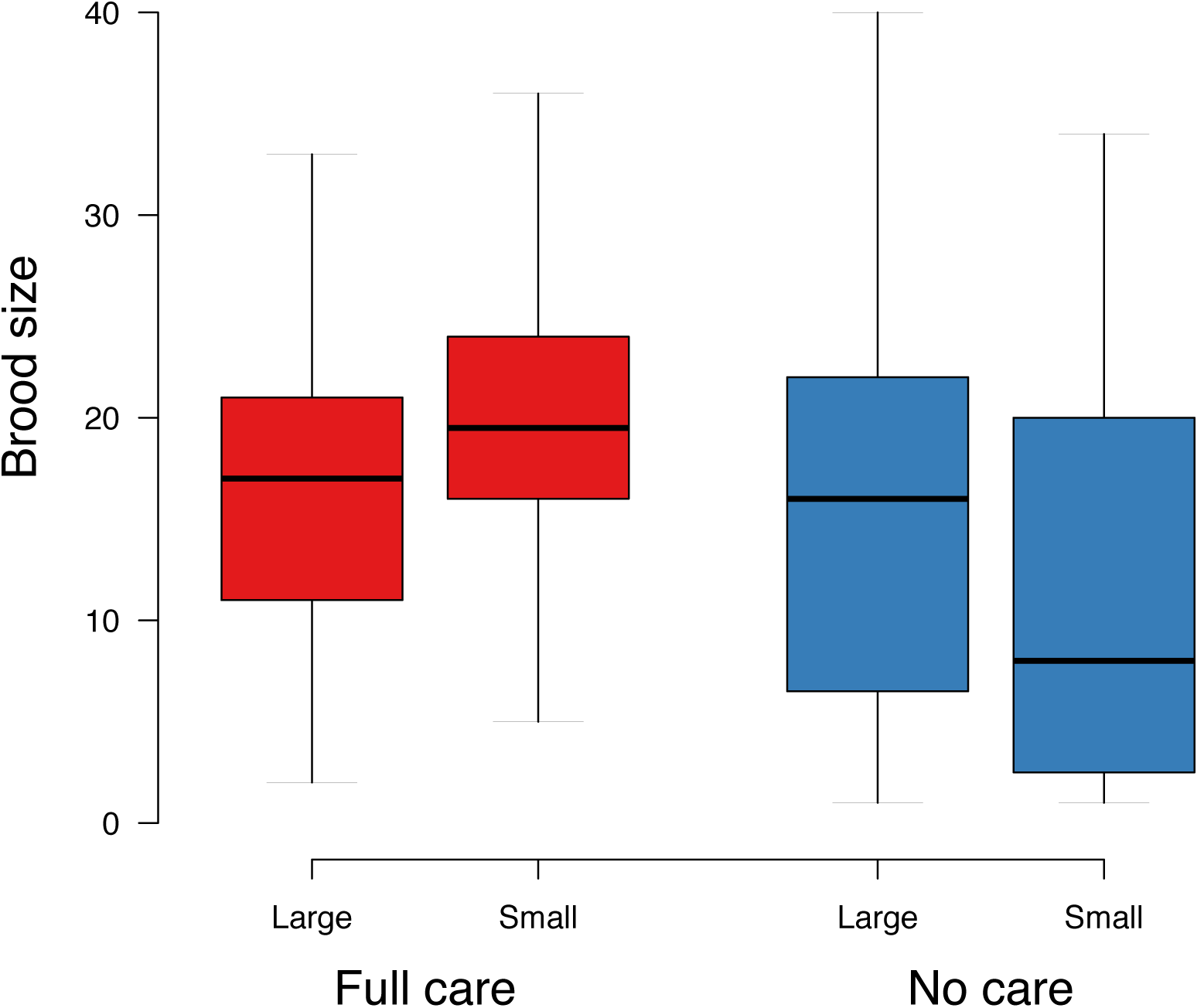
Brood size at larval dispersal in the four different experimental treatments in the selection experiment: Full Care Large (N=54) and Full Care Small (N=52) (in red); No Care Large (N=47) and No Care Small (N=15) (in blue). Both replicates per treatment were combined. Box plots show the median and interquartile ranges of the data.

## Tables

**Supplementary Table S1.**
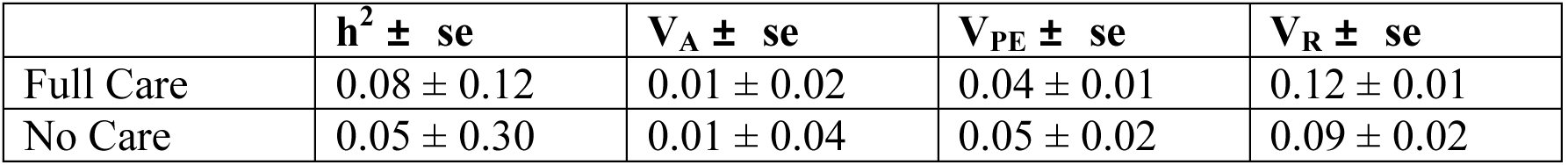
The variance components for pronotum width in the Full Care and No Care environments.

**Supplementary Table 2.** The realised heritabilities, and the associated standard errors, of adult body size for each of the eight experimental populations (i.e. the four experimental treatments, each replicated twice). For each population, the slope of the regression of cumulative response to selection against cumulative selection differential of each population was tested against zero. The t-values and P-values give the results of these tests.

| Population | Realised heritability | Standard error | t value | P value |
| --- | --- | --- | --- | --- |
| Full Care Large 1 | 0.048 | 0.025 | 1.929 | 0.112 |
| Full Care Large 2 | 0.130 | 0.024 | 5.401 | 0.003 |
| Full Care Small 1 | -0.042 | 0.041 | -1.030 | 0.350 |
| Full Care Small 2 | 0.022 | 0.019 | 1.140 | 0.306 |
| No Care Large 1 | 0.067 | 0.064 | 1.047 | 0.343 |
| No Care Large 2 | -0.023 | 0.024 | -0.917 | 0.401 |
| No Care Small 1 | -0.134 | 0.051 | -2.617 | 0.047 |
| No Care Small 2 | -0.076 | 0.043 | -1.771 | 0.137 |

**Supplementary Table 3.** The *Nicrophorus* species which were measured for size. The sample size (N) is the number of individuals photographed and measured from collections held in the Natural History Museum, London.

| Species | Number of photographs (N) | Mean pronotum width (mm) | Standard deviation |
| --- | --- | --- | --- |
| *Nicrophorus americanus* | 27 | 10.56 | 1.04 |
| *Nicrophorus antennatus* | 8 | 5.92 | 0.76 |
| *Nicrophorus apo* | 2 | 5.20 | 0.55 |
| *Nicrophorus argutor* | 5 | 6.67 | 0.42 |
| *Nicrophorus carolinus* | 40 | 6.99 | 0.93 |
| *Nicrophorus charon* | 9 | 5.52 | 0.65 |
| *Nicrophorus concolor* | 37 | 10.78 | 0.99 |
| *Nicrophorus dauricus* | 6 | 6.77 | 0.52 |
| *Nicrophorus defodiens* | 60 | 5.43 | 0.59 |
| *Nicrophorus didymus* | 28 | 5.51 | 0.60 |
| *Nicrophorus distinctus* | 14 | 6.73 | 0.46 |
| *Nicrophorus encaustus* | 5 | 5.67 | 0.50 |
| *Nicrophorus germanicus* | 24 | 9.92 | 1.38 |
| *Nicrophorus guttula* | 50 | 5.72 | 0.73 |
| *Nicrophorus heurni* | 12 | 5.35 | 0.55 |
| *Nicrophorus humator* | 33 | 7.16 | 0.85 |
| *Nicrophorus hybridus* | 8 | 7.26 | 0.89 |
| *Nicrophorus insularis* | 8 | 5.87 | 0.48 |
| *Nicrophorus interruptus* | 35 | 5.81 | 0.60 |
| *Nicrophorus investigator* | 105 | 5.99 | 0.77 |
| *Nicrophorus japonicus* | 16 | 6.58 | 0.98 |
| *Nicrophorus kieticus* | 29 | 4.34 | 0.53 |
| *Nicrophorus lunatus* | 5 | 6.65 | 1.10 |
| *Nicrophorus maculifrons* | 12 | 5.72 | 0.91 |
| *Nicrophorus marginatus* | 69 | 6.38 | 0.96 |
| *Nicrophorus mexicanus* | 20 | 5.84 | 0.76 |
| *Nicrophorus montivagus* | 18 | 4.32 | 0.53 |
| *Nicrophorus morio* | 7 | 9.00 | 0.88 |
| *Nicrophorus nepalensis* | 90 | 5.22 | 0.58 |
| *Nicrophorus nigricornis* | 4 | 6.34 | 1.00 |
| *Nicrophorus nigrita* | 15 | 6.07 | 0.99 |
| *Nicrophorus oberthuri* | 9 | 5.51 | 0.60 |
| *Nicrophorus obscurus* | 36 | 6.97 | 0.99 |
| *Nicrophorus olidus* | 20 | 4.66 | 0.62 |
| *Nicrophorus orbicollis* | 39 | 6.65 | 0.82 |
| *Nicrophorus podagricus* | 80 | 6.10 | 0.62 |
| *Nicrophorus przewalskii* | 4 | 6.59 | 0.20 |
| *Nicrophorus pustulatus* | 18 | 7.05 | 0.93 |
| *Nicrophorus quadrimaculatus* | 4 | 4.92 | 0.76 |
| *Nicrophorus quadripunctatus* | 56 | 5.05 | 0.67 |
| *Nicrophorus sayi* | 40 | 6.12 | 0.65 |
| *Nicrophorus scrutator* | 7 | 5.94 | 1.50 |
| *Nicrophorus semenowi* | 3 | 5.18 | 0.90 |
| *Nicrophorus sepultor* | 12 | 6.12 | 0.54 |
| *Nicrophorus smefarka* | 4 | 4.13 | 0.42 |
| *Nicrophorus tenuipes* | 20 | 5.63 | 0.39 |
| *Nicrophorus tomentosus* | 50 | 5.46 | 0.59 |
| *Nicrophorus vespillo* | 50 | 5.72 | 0.77 |
| *Nicrophorus vespilloides* | 70 | 4.83 | 0.59 |
| *Ptomascopus morio* | 23 | 4.16 | 0.56 |

**Supplementary Table 4.** Variation in the provision of parental care across burying beetle species. ‘Obligate’ care means that larvae cannot survive to their third instar unless they are cared for by their parents; ‘facultative’ care means larvae can survive without their parents.

| Species | Parental care | Source of information |
| --- | --- | --- |
| *Nicrophorus americanus* | Obligate | D. Howard pers. comm. |
| *Nicrophorus defodiens* | Facultative | (30) |
| *Nicrophorus humator* | Obligate | BP Springett cited in (30) |
| *Nicrophorus investigator* | Obligate | BP Springett cited in (30) |
| *Nicrophorus marginatus* | Obligate | D. Howard pers. comm. |
| *Nicrophorus mexicanus* | Facultative | (48) |
| *Nicrophorus nepalensis* | Facultative | S.-J. Sun pers. Comm. |
| *Nicrophorus orbicollis* | Obligate | (30,49) |
| *Nicrophorus pustulatus* | Facultative | (30,49) |
| *Nicrophorus quadripunctatus* | Facultative | (50) |
| *Nicrophorus sayi* | Obligate | (30) |
| *Nicrophorus tomentosus* | Facultative | (30) |
| *Nicrophorus vespillo* | Facultative | (51), B.J.M. Jarrett, unpub data |
| *Nicrophorus vespilloides* | Facultative | (19,21,49) |

